# Portable In-Field DNA Sequencing for Rapid Detection of Pathogens and Antimicrobial Resistance: A Proof-of-Concept Study

**DOI:** 10.1101/2024.10.05.616767

**Authors:** Stephanie Pillay, David Calderón-Franco, Thomas Abeel

## Abstract

**Purpose:** Infectious diseases represent a major global health threat due to the rapid spread of pathogens, leading to widespread outbreaks. Concurrently, antimicrobial resistance (AMR) is increasing, making standard treatments less effective and complicating infection management. Effective surveillance systems are essential to address these challenges.

**Methods:** We conducted a proof-of-concept study to evaluate a portable in-field microbial lab against a traditional molecular lab for DNA isolation, sequencing, microbial detection, antibiotic resistance gene identification, and plasmid classification. Samples from lake water, wastewater treatment plant sludge, and retail meat were selected to reflect relevant surveillance vectors. This approach provides valuable data for environmental monitoring, public health, and food safety, aiding in outbreak preparedness. We compared results using five metrics: DNA yield and purity, read N50, taxonomic classification, antibiotic resistance gene identification (ARGs), and plasmid classification.

**Results:** Our study found that metagenomic bacterial DNA isolation from environmental and food sources is feasible with portable lab technology, producing ≥800ng of DNA, suitable for Nanopore sequencing. DNA from retail meat, lake, and sludge samples resulted in similar read numbers and read N50 values. Taxonomic classification was achieved at the genus and species levels. A Jaccard similarity of over 50% was observed in the top 20 most abundant species between chicken samples, and lake samples. ESKAPE pathogens were detected in chicken and lake samples. ARGs and plasmids were also identified in both retail meat and lake samples.

**Conclusion:** This study demonstrates that in-field DNA isolation and sequencing are feasible with minimal equipment, yielding results comparable to traditional lab methods.

## 1 Introduction

Infectious diseases are one of the leading causes of death worldwide. Pathogens rapidly evolve and spread disease, leading to more virulent forms, including antimicrobial resistant bacteria [1]. The World Health Organisation (WHO) has named antimicrobial resistance (AMR) as one of the top ten global health threats among air pollution, non-communicable diseases, influenza and Ebola [2]. In 2019, infectious diseases caused an estimated 13.7 million deaths, including 4.95 million AMR-associated deaths. This number is projected to rise to 10 million deaths annually by 2050 [3, 4]. Organisations such as the Food and Agriculture Organization of the United Nations (FAO), the World Organisation for Animal Health (WOAH), and WHO have been working together to implement global action plans to tackle the burden caused by AMR [4]. This involves the strengthening of knowledge and evidence through research and surveillance systems as well as the monitoring of AMR across clinical and non-clinical sectors in all countries [5, 6].

Over the past decade, infectious diseases and AMR have been studied using traditional culture-dependent approaches and other molecular diagnostic techniques such as polymerase chain reaction (PCR), electrophoresis, multi-locus sequence typing (MLST) and pulsed-field gel electrophoresis (PFGE) [6–8]. These methods, while effective, provide limited information on only specific pathogens, can fail to detect pathogens, only detect culturable bacteria, are time-consuming, require trained personnel and dedicated laboratories and can be expensive [1, 6, 7, 9]. Unfortunately, due to the rapid spread and evolution of pathogens and microorganisms, new rapid detection techniques are needed [1, 7].

The COVID-19 pandemic highlighted the need for rapid detection methods as the surge in testing overwhelmed the local capacities in different countries [10]. Conventional diagnostic workflows for COVID-19 detection, relying on specialised equipment and supplies, faced shortages, creating an urgent need for cost-effective solutions [10–13].

In response, platforms utilising Third Generation Sequencing (TGS) were developed to facilitate real-time surveillance of viral pathogens. Examples of such platforms employed long-read Oxford Nanopore Technologies (ONT) sequencing, integrated with Isothermal Rapid Viral Amplification for Near Real-time Analysis (NIRVANA), CalmBelt and sequence variation analysis to facilitate rapid COVID-19 genome characterisation for outbreak tracking and diagnostics [12, 14, 15]. Such technologies offer the added benefit of portability, making them suitable for deployment in various locations, including wastewater monitoring [12, 16]. Additionally, a study comparing ONT sequencing with Illumina sequencing for COVID-19 detection found that ONT was not only faster but also more cost-effective, reducing both sequencing time and costs. This efficiency was further enhanced by using the rapid sequencing kit (SQK-RBK), which significantly reduced library preparation time [17].

The utility of ONT sequencing extends beyond COVID-19, proving valuable in antimicrobial resistance research. For example, a study by Martin et al. [18], utilised 16S rDNA amplicon sequencing with the ONT MinION and a portable Bento Bio Pro lab for detecting antibiotic-resistant *Campylobacter* in chickens in less than 5 hours. Similarly, Serpa et al. [19] employed ONT sequencing for detecting bacteria causing lower respiratory infections using CRISPR/cas9 techniques. ONT metagenomic sequencing has also demonstrated its capability to identify, characterise, and trace pathogens across diverse environments, including clinical, agricultural, environmental, and food safety contexts [9, 18–22]. As discussed by Bloemen et al. [9], ONT sequencing has been successfully applied in remote locations such as the ice caps in Iceland, sea ice in Allen Bay, Canada, and desert areas in Spain. These studies, utilising portable laboratory equipment and the ONT MinION device, have demonstrated that in-field sample processing can be performed effectively without issues related to sample transportation or preservation, thus ensuring reliable taxonomic classification. Despite its advantages, the ONT MinION sequencer, though portable, still requires pre-sequencing procedures, such as DNA isolation and library preparation, to be conducted in a traditional laboratory setting. The development of portable devices like the Bento lab [23] offers a solution by enabling genomics experiments to be conducted in various locations without the need for the resources typically found in molecular labs. Such equipment is cost-effective as it consists of a thermocycler, heating block, gel electrophoresis, transilluminator and centrifuge. It is also available in a cheaper, entry-level model and the more expensive pro-level model, both of which can be connected to a battery to conduct in-field genomics experiments [24–26]. Additionally, the Flongle adaptor can be used with the MinION sequencing device. While it does enable the use of lower throughput, the Flongle flow cells may offer a cost-effective approach to long-read sequencing which has been suitable for bacterial identification in clinical environments [27]. Such a portable laboratory device combined with a MinION and Flongle adaptor can be useful for antimicrobial resistance research, surveillance and in-field microbial detection.

In this proof-of-concept study, we first developed a modified DNA isolation protocol that can be used with the portable Bento lab (Pro-level) to yield high-quality DNA. Secondly, we assessed the differences between DNA isolated using a Bento lab (BL) and a traditional molecular lab (TL). We evaluated differences based on the following criteria: (i) DNA yield and purity, (ii) total nucleotide volume generated (Kb) and read N50 (Kb), (iii) similarity in taxonomic composition, (iv) identification of antibiotic resistance genes (ARG) and (v) classification of plasmids.

## 2 Methods and materials

### 2.1 Sample collection and processing

Three different samples were collected to assess their environmental and food safety impacts; lake water (n=1), sludge (n=1) and retail meat product (n=1). 2L of lake water was collected in a sterile bottle from the surface (5cm - 10cm) of a recreational lake in Delft, The Netherlands (52°01’18.2”N 4°22’59.3”E). Retail poultry (whole carcass) was purchased from a supermarket in Delft, The Netherlands and washed with 1L of distilled water. The carcass rinsate from the retail poultry bag was transferred into a sterile bottle for further processing. Lastly, 1.8mL of sludge was obtained at a wastewater treatment plant in Amsterdam West, The Netherlands. All samples were collected and stored on ice until DNA isolation was done on the same day.

### 2.2 DNA isolation

To assess the differences between the portable pro-level Bento lab (BL) (Bento Bioworks, UK) and a traditional molecular lab (TL), DNA was isolated from all samples using both these lab types (n=6) and concentration was assessed using a Qubit 3 (Thermo Fisher Scientific, Wilmington, DE, USA). DNA was isolated from the sludge and carcass rinsate samples using the DNeasy Ultraclean Microbial extraction kit (Qiagen Inc., Valencia, CA) and from the lake water using the DNeasy PowerWater DNA kit (Qiagen Inc., Valencia, CA). Before isolation, 300mL of lake water was filtered using a vacuum pump with a 0.2mm cellulose filter (Merck Millipore, Ireland). DNA isolation was done following the manufacturers’ instructions in the traditional molecular lab.

With regards to the Bento lab, we chose the pro-level model which has a varying centrifugation speed (max 13 500 RPM). The centrifuge capacity is also limited to 2ml Eppendorf tubes which can be overcome by increasing centrifugation times during experiments. Modifications to the DNA isolation protocols were made for the Bento lab as follows:

#### Modifications to the DNeasy PowerWater DNA extraction kit

Step 8: Transfer the supernatant to a clean 2ml collection tube *→* transfer to 2ml

Eppendorf tube.

Step 9: 13 000 x g for 1 min *→* 8000 x g for 3 min.

Step 10: Transfer the supernatant to a 2ml collection tube *→* transfer to 2ml Eppendorf tube.

Step 12: 13 000 x g for 1 min *→* 8000 x g for 3 min.

Step 13: Transfer the supernatant to a 2ml collection tube *→* transfer to 2ml Eppendorf tube.

Step 15-20: Load 650*µ*l of supernatant onto an MB spin column *→* place MB spin column in a 1.5ml Eppendorf tube with cap cut off (modified collection tube). Centrifuge at 13 000 x g for 1 min *→* 8000 x g for 3 min.

Step 22: 13 000 x g for 1 min *→* 8000 x g for 3 min.

#### Modifications to the DNeasy Ultraclean Microbial kit

Step 1: Add 1.8ml of microbial (bacteria, yeast) culture to a 2ml collection tube *→*

1.8ml carcass rinsate / sludge in a 2ml Eppendorf tube. Centrifuge at 10 000 x g for 1 min *→* 8000 x g for 2 min. Decant the supernatant.

Centrifuge at 10 000 x g for 1 min *→* 8000 x g for 2 min.

Step 2: Transfer resuspended cells to a PowerBead Tube *→* PowerBeads from the PowerBead Tube were added to a 2ml Eppendorf tube.

Step 5: Centrifuge the tubes at a maximum of 10 000 x g for 30 s *→* 8000 x g for 2 min, Step 6: Transfer the supernatant to a clean 2 ml Collection Tube *→* transfer to 2ml Eppendorf tube.

Step 8: Centrifuge the tubes at 10 000 x g for 1 min *→* 8000 x g for 3 min.

Step 9: Transfer the entire volume of supernatant to a clean 2 ml Collection Tube *→*transfer to 2ml Eppendorf tube.

Step 11: Load 700*µ*l of supernatant onto an MB spin column *→* place MB spin column in a 1.5ml Eppendorf tube with cap cut off (modified collection tube) and load 700*µ*l of supernatant to spin column.

Centrifuge at 10 000 x g for 30s *→* 8000 x g for 3 min.

Discard the flowthrough and add the remaining supernatant to the MB Spin Column *→* place the MB spin column in a 1.5ml Eppendorf tube with cap cut off (modified collection tube).

Centrifuge again at 10 000 x g for 30s *→* 8000 x g for 3 min.

Step 12: Add 300*µ*l of Solution CB and centrifuge at 10 000 x g for 30s *→* 8000 x g for 3 min.

Step 13: Discard the flow-through. Centrifuge at 10 000 x g for 1 min *→* 8000 x g for 3 min.

Step 14: Place the MB Spin Column in a new 2 ml Collection Tube *→* place MB spin column in a new 1.5ml Eppendorf tube with cap cut off (modified collection tube). Step 16: Centrifuge at 10 000 x g for 30s *→* 8000 x g for 3 min.

Step 17: Discard the MB Spin Column. The DNA is now ready for downstream applications *→* Store the flowthrough in a 1.5ml or 2ml Eppendorf tube.

### 2.3 ONT Nanopore Sequencing

DNA libraries were prepared using only the Bento lab and the Rapid Sequencing Kit V.14 (SQK RAD114) (Oxford Nanopore Technologies, USA) as per the manufacturer’s instructions for the Flongle application with Flongle flow cells (R10.4.1. FLO-MIN114). 75ng of DNA isolated using both lab types (n=6) and the Zymobiomics microbial community standard was used as a control (n=1) (Zymo Research Corporation, Irvine, CA, USA). Sequencing of all samples was run for 24 hours. The MinKNOW GUI (ONT, v.19.05.0) was used for setting appropriate sequencing parameters (analysis protocol, flow cell type, run duration, basecalling and for following the progression of the sequencing runs in real-time (number of reads, read length distribution and read quality, and pore activity status).

### 2.4 Bioinformatic analysis

Raw .Pod5 files were basecalled using Dorado v.0.4.3 with the (–model dna_r10.4.1_e8.2_400bps_sup@v4.2.0). Basecalled bam files were converted to FASTQ files using Samtools v.1.9 and further processed using NanoFilt v.3 [28]. Taxonomic classification on the filtered reads representing each sample (n=7) was done using a modified protocol provided by (Taxonomic-Profiling-Minimap-Megan) at:(https://github.com/PacificBiosciences/pb-metagenomics-tools). The NCBI nt database was downloaded (11-06-2024) and indexed according to Portik et al., [29].

Minimap2 v.2.26 [30] was used to align the raw reads to the NCBI nt database using the -ac map-ont –sam-hit-only –secondary=no –split-prefix command to generate a sam file. Sam files were converted to rma files using MEGAN v.6.25.9 [31] with the sam2rma function, megan-nucl-Feb2022.db and -lg -ram option to generate rma files with assigned read counts. The MEGAN6 community edition v.6.19 was used to visualise the rma files and export the taxonomic classifications with assigned read counts.

Antibiotic resistance genes were detected using BLAST v.2.11.0 [32] against the Comprehensive Antibiotic Resistance Database (CARD) v.4.0.2 [33] with an 80% identity and 60% query coverage to obtain the number of antibiotic resistance genes identified, the antibiotic resistance gene families and corresponding drug class. Reads were classified as plasmids by Plasclass v.0.1.1 [34] and were considered plasmid originating when a probability of over 0.75 was assigned to a read. Plasmid-assigned reads were cross-referenced with the identified antibiotic resistance genes to ascertain the presence of antibiotic resistance genes within the plasmids.

## 3 Results

In this proof-of-concept study, we first aimed to determine if a modified DNA isolation protocol can be used with a portable lab (Bento lab) to yield high-quality DNA. Secondly, we compared the results generated from using a Bento lab to a traditional laboratory across various metrics: reads generated, read N50, total data produced, microbial classification, detection of antibiotic resistance genes, and plasmid classification.

### 3.1 DNA isolation is feasible with modified protocol and Bento lab

To assess the differences between the DNA isolated from different samples obtained by the Bento lab (BL) and the traditional lab (TL), we measured the DNA yield and purity. The prolevel Bento lab faces limitations due to its centrifugation speed and the size of Eppendorf tubes that can be used, which are critical factors in DNA isolation protocols. To address these constraints, a modified protocol is required that can still deliver high-quality DNA with good purity. DNA yield and purity are crucial in sequencing as they can directly impact the quality of the results. A high DNA yield ensures that there is enough DNA to be sequenced while purity ensures that no contaminants are affecting the sequencing process [35].

We isolated metagenomic bacterial DNA with sufficient yield from each sample using both the modified DNA isolation protocol with the Bento lab and the unmodified protocol with the traditional lab. We measured the DNA yield and purity of each DNA sample isolated from the Bento lab (n=3) and the traditional lab (n=3). Table 1 shows the DNA yield (ng) isolated from the lake, sludge and the retail meat product. The DNA yield was sufficient for downstream sequencing for all sample types. The purity of the DNA samples varied, with measurements taken at A260/A280. As indicated in Table 1, all samples, excluding Lake_BL, had a DNA purity ranging from 1.8 - 1.9 which can be considered pure according to past metagenomic research [35–37]. Interestingly, the DNA isolated from the sludge and the retail meat product had similar values of 1.9 which could be attributed to the use of the same DNA isolation kit. However, we expected to see differences between the Bento lab samples and the traditional lab samples due to the modifications in the DNA isolation protocol.

**Table 1:** Table showing the DNA yield (ng) and DNA purity (A260/A280) in the Chicken, Lake, and Sludge DNA samples isolated using the Bento lab and traditional lab.

| DNA yield (ng) produced |  |  |
| --- | --- | --- |
| Sample type | Bento lab (BL) | Traditional lab (TL) |
| Chicken | 800 | 1780 |
| Lake | 2340 | 1630 |
| Sludge | 11750 | 7580 |
| Purity of the DNA (A260/A280) samples |  |  |
| Sample type | Bento lab (BL) | Traditional lab (TL) |
| Chicken | 1.990 | 1.976 |
| Lake | 1.628 | 1.838 |
| Sludge | 1.900 | 1.897 |

The purity recorded for Lake_BL was 1.6, indicating the presence of phenol, protein, or other contaminants [38, 39]. This suggests that the modified DNA isolation protocol (DNeasy PowerWater DNA extraction kit) and the use of the Bento lab for the lake sample results in lower purity compared to the unmodified protocol used in the traditional lab. The Bento lab is restricted by centrifugation speed and the size of Eppendorf tubes that can fit into the centrifuge rotor. The lower purity suggests that these restrictions may impact the purity of DNA. Nevertheless, it has been stated that low phenol contamination in DNA samples has minimal effects on the rapid sequencing kit during the library preparation step and has no impact on the sequencing process [40, 41].

### 3.2 Sequencing samples isolated using the Bento lab and traditional lab yields similar results

To compare the differences between the reads generated, read N50 and total data produced between metagenomic DNA samples obtained using the Bento lab and the traditional lab, we sequenced each sample using the MinION with the Flongle adaptor. The number of reads generated during sequencing can influence taxonomic classification and the identification of ARGs and plasmids as a higher number of reads will lead to more accurate classifications [42]. A full summary can be found in Supplementary table S1.

Table 2 shows that the reads generated, read N50, and data volume produced from the same sample types isolated using both lab technologies were within a similar range. Firstly, the Zymobiomics microbial community was used as a control to ensure that the rapid sequencing kit and the MinION with the Flongle worked correctly, as library preparation was done only using the Bento lab. According to Oxford Nanopore Technologies, the Flongle can produce up to 2.6Gb data [43]. We produced 46Kb reads and 2.7Gb of data from sequencing the control.

**Table 2:** Summary of MinION sequencing results for metagenomic bacterial DNA isolated from control, lake, sludge, and chicken samples using both the Bento lab (BL) and traditional lab (TL).

| Sample type | Reads generated (K) | Read N50 (Kb) | Total data produced (Mb) |
| --- | --- | --- | --- |
| Control | 46.57 | 5.94 | 2780 |
| Chicken BL | 16.950 | 9.84 | 265.93 |
| Chicken TL | 14.432 | 7.24 | 230 |
| Lake BL | 21.515 | 11.8 | 964.35 |
| Lake TL | 20.111 | 5.63 | 1330 |
| Sludge BL | 5.678 | 102.42 | 93.75 |
| Sludge TL | 6.823 | 46.67 | 188.43 |

According to Table 2, the same sample types, regardless of their DNA isolation protocol or technology, generated a similar number of reads. While the rapid sequencing kit suggests a DNA input of 100ng - 150ng, we used 75ng of DNA during the library preparation step. This lower amount was chosen to optimise resource use and minimise costs when working with limited amounts of sample in the field. Additionally, using 75ng allowed us to test the sensitivity of the sequencing protocol and assess its performance with reduced input, ensuring that the library preparation and sequencing could still yield high-quality data. Unfortunately, the amount of DNA used in library preparation does affects the data that is produced as a lower amount of DNA will yield low data output and shorter length reads which is evident in our study. The rapid sequencing kit, while requiring fewer manipulations in the library preparation steps, does not include a clean-up step. This can impact the sequencing results, as residual contaminants may remain in the sample [44].

We hypothesised that the varying pore availability and differences in DNA fragmentation could impact the read count and read N50 values observed during sequencing. While the Flongle flow cell has a maximum of 126 channels, the manufacturer guarantees 60 available pores for sequencing [43]. In our study, we had 40 to 70 pores available during the sequencing runs of all samples (Supplementary table S1). While the lake and retail meat product samples generated a high number of reads, the DNA isolated from the sludge samples yielded a low number of much longer reads (5K reads, read N50 of 46Kb, 90Mb data). Due to the length of the DNA being sequenced, the pores within the flow cell can become blocked or lost i.e., inactive and stop operating, resulting in a low read count with a high read N50 [43]. We propose that a higher number of reads with a lower read N50 can be achieved by increasing the fragmentation time in the rapid sequencing kit protocol [45]. Currently, the fragmentation step (step 6) in the rapid sequencing kit states incubation should be done at 30*°*C for 1 minute and then 80*°*C for 1 minute however, we propose only increasing the incubation at 80*°*C to 2-4 minutes. Further analysis should be done on the effects of the fragmentation time on the read N50 and the reads generation by sequentially increasing the time at 80*°*C and observing the read N50 and reads generated during sequencing. Similar to our study, Maguire et al., [42] stated that the rapid sequencing kit, similar to the one used in our study, results in poor Nanopore sequencing reactions and low output of reads (0.3–1.7M reads) in agricultural water. However, a low number of reads can be sufficient for genus and species-level identification.

### 3.3 A high degree of similarity is seen in the top 20 most abundant species between Bento lab samples and traditional lab samples

To identify the microbial community at genus and species-level between the DNA isolated from the lake, sludge and retail meat product processed by the Bento lab and traditional lab, we performed taxonomic classification. Furthermore, we assessed similarities between the microbial communities found in the Bento lab and traditional lab samples. Details on the relative abundances of the bacterial genera, species, common bacterial genera and species and similarity are in Supplementary tables S2 -S5.

Species with a higher number of assigned reads were common between the same sample types regardless of the lab technologies. Firstly, food spoilage bacteria were present in both retail meat product (chicken) samples. The *Pseudomonas* genus was dominant in both chicken samples each of which was isolated using both lab technologies (Supplementary table S2). On a genus level, we identified the same number of genera in the DNA isolated using both lab types (Figure 1). Only 8 of these

**Fig 1:**
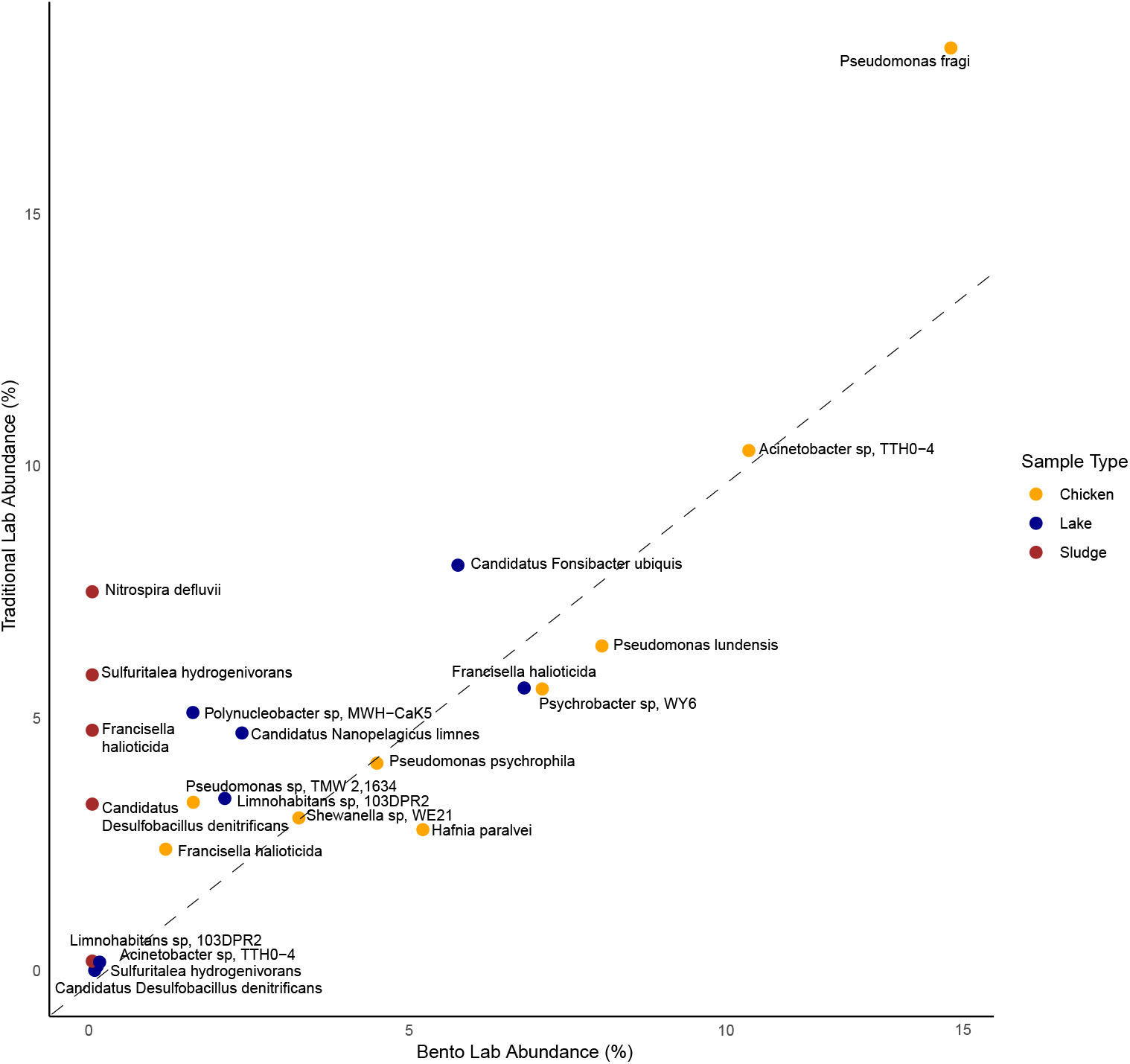
The abundances of bacterial species present in the chicken, lake and sludge samples processed by the Bento lab and the traditional lab. The x-axis represents the abundances of bacterial species detecting from Bento lab samples and the y-axis represents the abundances of bacterial species detected from the traditional lab samples. All bacterial species shown in this figure have abundances of over 3%. Dots in the figure are bacterial species found within a specific sample type and are colour coded accordingly (Chicken = orange, Lake = blue and Sludge = brown.)

genera (*≥* 1%) were common between the Chicken_BL and Chicken_TL, resulting 10 in a Jaccard similarity index of 0.43 (Supplementary tables S4 & S5), indicating a moderate similarity in the bacterial communities identified between the two chicken samples. The Jaccard similarity index increased to 0.54 when focusing on the top 20 most abundant genera, suggesting a higher degree of similarity in the most dominant bacteria identified between the two samples (Supplementary table S5). Figure 1 shows the species-level identification revealing that *Pseudomonas fragi* was the dominant species present in both Chicken_BL and Chicken_TL (Supplementary table S3). *Pseudomonas fragi* is commonly found on retail meat products and contributes to the spoilage of meat which can potentially be a health hazard [46, 47]. In total, 237 bacterial species were identified between the chicken samples however, only 19 species were common (*≥* 1%) and had a Jaccard similarity of 0.4449 (Supplementary tables S4 & S5). This similarity increased to 0.9047 when focusing on the top 20 abundant species, indicating a high level of consistency in identifying the most abundant species between the two samples regardless of the adapted DNA protocol.

Figure 1 shows environmental freshwater bacteria were detected in both lake samples. The top three genera identified between lake samples were the *Flavobacterium, Limnohabitans* and *Polynucleobacter* bacteria (Supplementary table S2). While a higher number of genera were detected in Lake_BL (413) compared to Lake_TL (276), 15 similar bacterial genera (*≥* 1%) were detected between the two lake samples (Supplementary table S2). A Jaccard similarity of 42% was calculated indicating that only 42% of the bacterial genera identified are common between Lake_BL and Lake_TL (Supplementary table S5). Similar to the chicken samples, the similarity increased when only focussing on the top 20 abundant genera (66%). On the species level, we identified a total of 1039 different species with 11 species in common between the lake samples (*≥* 1%) (Supplementary tables S3 & S4). The Jaccard similarity index of the top 20 abundant species in Lake_BL and Lake_TL was 0.666 indicating a moderate level of overlap between the two samples, with approximately 66.6% of the top 20 abundant species being common to both lake samples (Supplementary table S5).

The Jaccard similarity may be influenced by the number of reads assigned to taxa during the taxonomic classification step. In this study, approximately 8000 reads in the Lake_BL sample were classified compared to 1900 reads in the Lake_TL sample. These differences indicate that more bacterial species were detected in the Lake_BL sample, contributing to a greater number of unique genera and, consequently, a lower overlap between the two samples (Supplementary table S1).

Regarding the DNA isolated from sludge, we could only perform taxonomic classification on the Sludge_TL sample as shown in Figure 1 (Supplementary table S3). This was due to the insufficient number of reads generated when the Sludge_BL was sequenced (Table 2). We identified a total of 159 genera and 266 species in Sludge_TL with the predominant species being *Nitrospira defluvii* which is commonly detected in sludge (Supplementary tables S2 & S3) [48–50].

Overall, our study demonstrates that while there are differences in the observed microbial communities between the DNA isolated using the Bento lab and the traditional lab, there is a moderate to high level of similarity between the most abundant species. This suggests that both methods are effective for microbial community detection, although the use of traditional labs with no modified DNA isolation protocol may

yield more comprehensive results in some cases, such as with sludge samples. Additionally, the use of the Flongle can enhance the capabilities of portable labs like the Bento lab for in-field microbial detection. However, the reduced data volume of the Flongle provides a more limited view of the microbial community, primarily detecting the most abundant species compared to the MinION [47, 51]. While this provides valuable insight into the dominant microbial populations in environmental and food sources, it may not capture the full diversity of less abundant species present in the microbial community.

### 3.4 ESKAPE pathogens can be detected in low abundances

To determine the presence of the ESKAPE (*Enterococcus faecium, Staphylococcus aureus, Klebsiella pneumoniae, Acinetobacter baumannii, Pseudomonas aeruginosa*, and *Enterobacter* spp.) pathogens in the lake, sludge and retail meat product (chicken) samples and assess the similarities between samples processed using the Bento lab (BL)and the traditional lab (TL), we investigated the reads assigned during taxonomic classification.

Figure 2 shows the presence of ESKAPE pathogens in the lake, sludge and chicken DNA samples. We identified *Pseudomonas aeruginosa* and *Acinetobacter baumanni* in both the chicken samples (*≤*1%) with *Klebsiella pneumonia* and *Enterobacter* sp. detected in the Chicken_BL (*≤*1%) (Supplementary table S6). Additionally, we detected *Pseudomonas aeruginosa* and *Enterobacter* in the lake samples (Supplementary table S6). The presence of ESKAPE pathogens in the lake and chicken samples may be concerning. However, ESKAPE pathogens are not generally pathogenic, as they are natural colonisers of humans and animals [52, 53]. Additionally, these microorganisms can be commonly found in soil and aquatic environments [54, 55]. Further analysis on confirming pathogenic strains should be done either via conducting polymerase chain reaction (PCR), mass spectrometry (MS), Matrix-assisted laser desorption ionization-time-of-flight mass spectrometry (MALDI-TOF MS) or via the use of computational tools such as StrainGE [56, 57].

**Fig 2:**
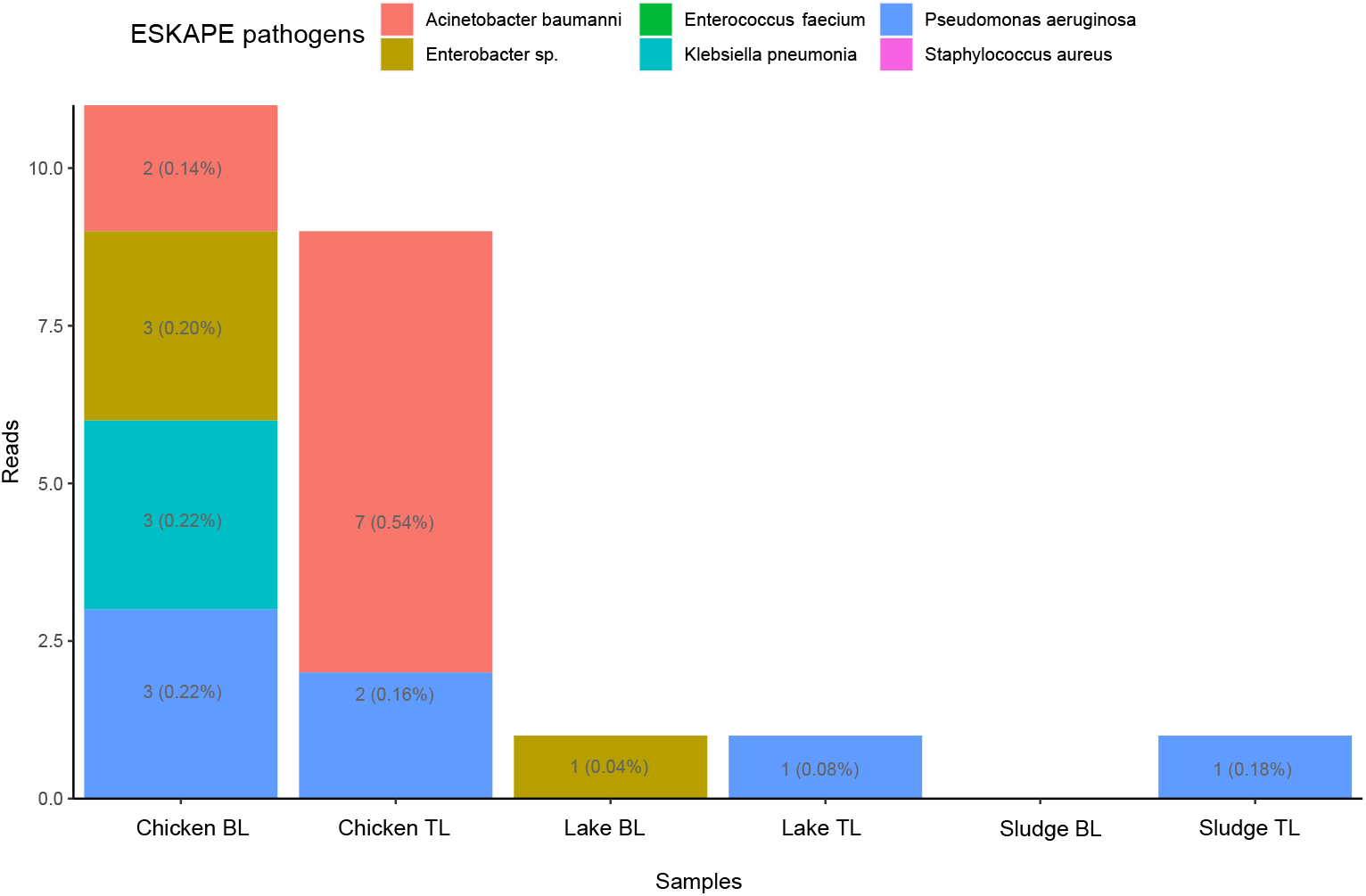
The number of reads assigned to ESKAPE pathogens detected in each DNA sample isolated using the Bento lab and the traditional lab. The x-axis indicates the different samples and the y-axis indicates the number of reads assigned to the ESKAPE pathogen. The number of reads and corresponding abundances in percentages for each ESKAPE pathogen are indicated in text within each bar. Each ESKAPE pathogen is indicated in different colours.

Lastly, due to the lack of data in Sludge_BL, we were unable to detect ESKAPE pathogens however, we did detect *Enterococcus faecalis* and *Pseudomonas aeruginosa* in Sludge_TL. It is evident by the read count that there is difficulty in the detection of pathogens and other bacteria as after the pre-processing steps, the Sludge_BL sample had only 184 reads compared to the 10K reads of Sludge_TL.

The variation in the reads assigned to the ESKAPE pathogens between samples processed by the Bento lab and the traditional lab can largely be attributed to differences in read counts generated during the sequencing run. Higher read counts typically result in more comprehensive and accurate microbial identification. This variation highlights the importance of optimising the pre-sequencing and sequencing protocols to ensure consistent and reliable detection of microbial communities, particularly for environmental and food safety applications.

### 3.5 Family level similarity for AMR genes

To determine the similarities between antibiotic resistance genes (ARGs) in the lake, sludge and retail meat product (chicken) DNA samples that were isolated using the Bento lab and traditional lab, we investigated the presence of ARGs. Identifying the presence of ARGs is crucial for public health as bacteria carrying these genes can potentially cause infections. Additionally, the identification of ARGs in environmental and food sources can assist in mitigating the spread of AMR to other environmental and clinical sectors [6].

Figure 3 shows the identification and number of ARGs with corresponding AMR gene families present in the different lake, sludge and chicken samples processed by the Bento lab and traditional lab. The RND efflux pump gene family was commonly detected. We identified 10 different ARGs in Chicken_BL and 11 different ARGs in Chicken_TL with only 3 ARGs in common (Supplementary table S7). However, 90% of the ARGs detected in both chicken samples were from the RND efflux gene pump family. On the gene level, 3 *MexF* genes were detected in Chicken_BL conferring resistance to cephalosporin, macrolides, aminoglycosides, and fluoroquinolone while 2 *YajC* and 2 AxyY genes were predominant in the Chicken_TL sample, conferring resistance to multiple antibiotic drug classes: tetracyclines, disinfecting agents and antiseptics, rifamycin, glycylcycline, cephalosporin, penam, phenicol, oxazolidinone, fluoroquinolone, glycopeptide, macrolide and aminoglycosides (Supplementary table S7). This is similar to a study conducted by Heir et al., [58] who detected the RND efflux pump gene family in retail products from Norwegian slaughterhouses. The RND efflux pump gene family and the ARGs comprising the family are commonly associated with *Escherichia, Salmonella* and *Pseudomonas*, all of which were detected in our chicken samples [58–61].

**Fig 3:**
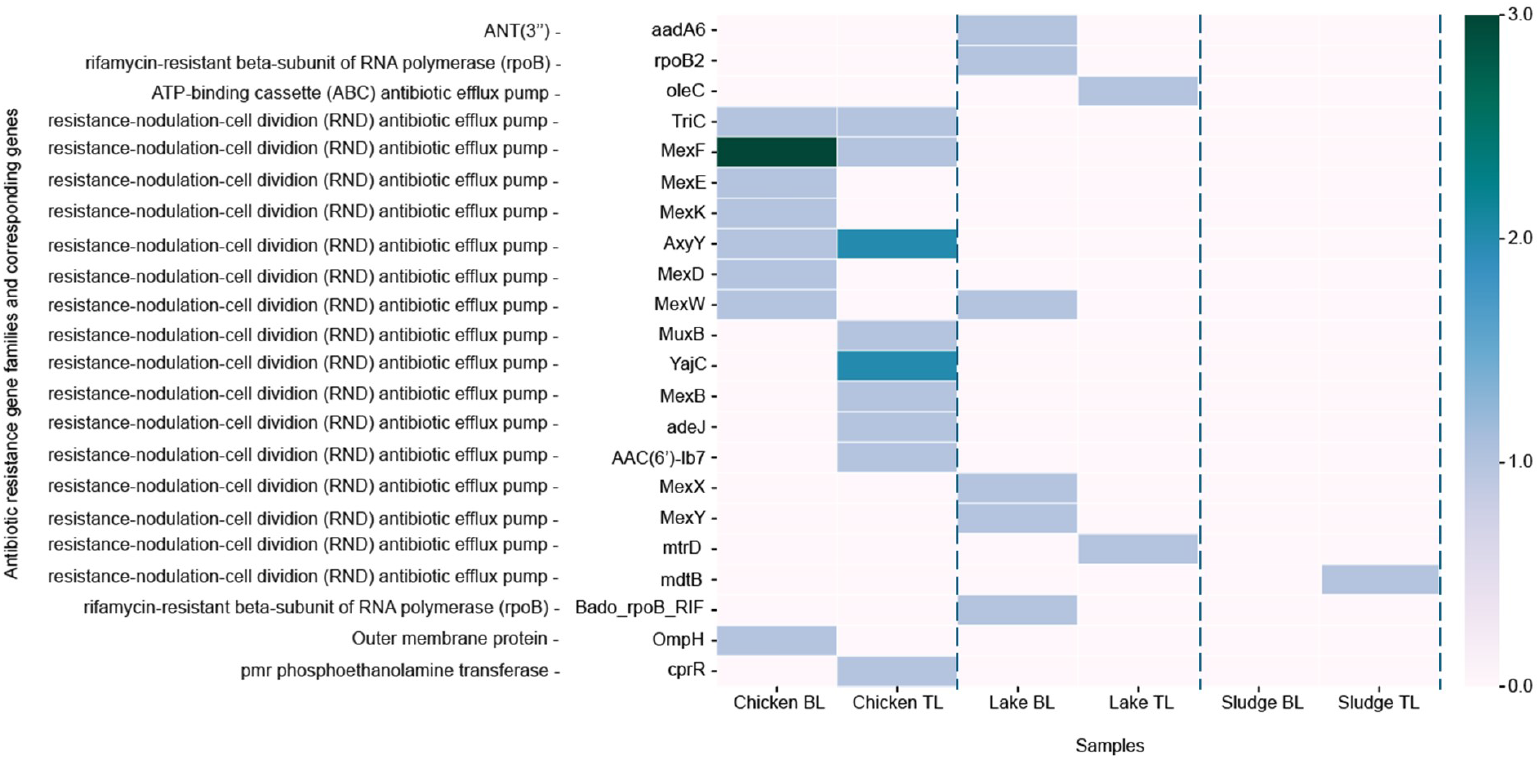
Heatmap showing the number of antibiotic resistance genes and corresponding gene families detected in each DNA sample isolated using the Bento lab and the traditional lab. The x-axis indicates the samples, the y-axis indicates the ARGs and AMR gene families and the colour intensity corresponds to the respective number of ARGs identified within their specific sample. The numbers are indicated by the colour intensity with low = pink and high = green and the dashed line indicates the separation between sample types

More variation in the detected ARGs was observed between the lake samples. Figure 3 shows that 50% of the ARGs identified in the Lake_BL and TL samples belong to the RND efflux pump gene family. We also detected ARGs conferring to aminoglycosides, rifamycin in Lake_BL and macrolides in Lake_TL with no common ARGs detected (Supplementary table S7). The detected AMR gene families align with the findings of Filipic et al. [62], who concluded that the RND efflux pump gene family was prevalent in a recreational lake. The detection of RND efflux pump gene families in both retail meat products and lake water may be a biosafety and food-safety hazard as these gene families can confer antibiotic resistance and multi-drug resistance in bacteria. They can be found in both pathogenic, non-pathogenic and opportunistic bacteria such as *Escherichia coli, Klebsiella pneumonia, Enterobacter spp, Pseudomonas aeruginosa, Acinetobacter baumanni* and emerging pathogens such as *Strenotrophomos maltophila* [63–65]. The RND efflux pump is naturally found in the chromosome of bacteria which can spread to humans and animals potentially

spreading antibiotic resistance and outbreaks.

While we see differences in the ARGs detected within similar sample types, there is a similarity in the AMR gene family level. Differences in the detection of ARGs can be attributed to the low number of reads or the percentage identity used during the alignment against the CARD database however, our study used an 80% identity, which previous studies have shown to be a strict cut-off [66]. Even though we did detect ARGs, the low number observed is likely due to the limited data volume generated, influenced by factors such as the DNA isolation process, library preparation, or the smaller output of the Flongle, which produces only 2.6Gb of data compared to the MinION’s 48Gb. Unfortunately, the number of reads to identify and confirm the presence of ARGs has not been defined yet as some studies have suggested one read is relevant due to the low threshold for long-read sequencing while others suggest that there should be at least 10 reads [67].

### 3.6 A higher number of plasmids detected in the chicken and lake bento lab samples

To determine the presence of plasmids between each sample type, we classified reads as plasmids. Furthermore, we investigated if the plasmids detected were carrying ARGs. Plasmids are important vehicles for rapid adaptation of bacterial populations changing to their environmental conditions. Plasmids can carry accessory genes such as ARGs and transfer these genes to pathogenic and non-pathogenic bacteria. An understanding of plasmids and the ARGs they carry is imperative in understanding the spread of antibiotic resistance and the role they play in emerging resistance and infections [68].

Table 3 shows the number of reads classified as plasmids in the metagenomic bacterial DNA isolated from the lake, chicken and sludge samples processed using the Bento lab and the traditional lab. We detected plasmids in all DNA samples isolated using the Bento lab and the traditional lab (Table 3). As expected the number of reads classified as plasmids within a sample type differed according to the number of reads within the dataset. A higher number of reads were generated from the Bento lab DNA samples, therefore a higher number of reads were classified as plasmid derived. The chicken samples had a difference of 9.16%, while the lake samples exhibited an 8.20% difference. In both instances, the Bento lab samples revealed a higher plasmid count. Interestingly, the Sludge_BL sample contained the lowest number of reads that could still be used to detect plasmids. This is likely because the reads in the Sludge_BL samples have longer read length (N50) which could make them more suitable for plasmid classification by PlasClass [34].

**Table 3:** The number of reads classified as plasmids and the number of reads associated with ARGs.

|  | Chicken<br>BL | Chicken<br>TL | Lake<br>BL | Lake<br>TL | Sludge<br>BL | Sludge<br>TL |
| --- | --- | --- | --- | --- | --- | --- |
| **Plasmids** | 544 | 496 | 2114 | 1944 | 8 | 324 |
| **Plasmids associated**<br>**with ARGs** | - | 1<br>AAC(6')-ib7 | 1<br>aadA6 | - | - | - |

Secondly, Table 3 shows two reads classified as plasmids that are associated with the previously detected ARGs. Both plasmids were associated with aminoglycoside resistance genes, *AAC(6’)-ib7* and *aadA6*, and detected in Chicken_TL and Lake_BL samples, respectively.

The differences in the number of plasmids identified and those associated with ARGs between the Bento lab and traditional lab samples may be attributed to differences in read numbers and read lengths. The higher number of reads from the Bento lab samples may have allowed for the detection of more plasmids, while the longer read lengths in the traditional lab samples may have facilitated the classification of plasmids even with fewer reads. This highlights the trade-offs between read numbers and read lengths in metagenomic analyses. As stated previously, the Flongle provides limited reads which may restrict the detection of plasmids especially those less abundant or present in low copy numbers.

## 4 Conclusion

Our proof-of-concept study has shown that the pre-sequencing steps can be achieved by using a portable pro-level Bento lab. We successfully isolated metagenomic bacterial DNA from environmental and food sources using a modified DNA isolation protocol and portable lab. We isolated DNA with sufficient yield needed for sequencing using both lab technologies and generated reads for each sample type using the MinION and Flongle.

We identified bacteria down to the species level from the DNA samples that were isolated using the Bento lab and the traditional lab. DNA isolated from the same samples showed bacterial similarities of over 40% however this increased between more abundant genera and species. We also identified ESKAPE pathogens in the lake, sludge and retail meat product (chicken) samples with an abundance of less than 1%. Furthermore, we detected ARGs in the lake, sludge and retail meat product (chicken) samples from both lab types. The RND efflux pump gene family was common between the same sample types. Lastly, we identified a similar number of plasmids in all samples with two plasmids associated with aminoglycoside resistance.

Since this study is a proof-of-concept, the study is limited by the number of samples and the limited amount of reads generated by the Flongle. To overcome this, future research should benchmark the modified DNA isolation protocol with the Bento lab and the unmodified DNA isolation protocol with the traditional molecular lab using a higher number of samples from various sources. This will aid in investigating how results from DNA concentration, purity, read counts, read N50 and identification differ between the Bento lab and the traditional lab. Furthermore, optimisation of presequencing steps using portable lab equipment such as the Bento lab needs to be conducted to improve DNA quality and data generation.

In this study, the Bento lab equipment was used during the library preparation step with the rapid sequencing kit which required fewer resources and less equipment, eliminating the need for modifications during the library preparation step. However, this streamlined process may not apply to other sequencing kits due to the limitations of the Bento lab setup. Additional experiments using various sequencing kits and adapting library preparation protocols with diverse samples to gather more comprehensive data, thereby supporting in-field research efforts is needed. Additionally, research into in-field microbial detection and AMR should also take into account and compare the data generated by the Flongle versus the MinION.

We believe that this proof-of-concept study can be the stepping stone into future research aiming at portability and real-time data generation essential for antibiotic resistance surveillance in different types of environments.

## Supplementary information

Supplementary tables can be found the Supplementary_materials.xlsx file.

## Declarations

### 4.0.3 Ethics approval and consent to participate

Not applicable

### 4.0.4 Consent for publication

All authors contributed to the article and approved the submitted version.

### 4.0.5 Data availability

The sequence data has been submitted to the Sequence Read Archive (SRA) under Bioproject PRJNA1168083.

### 4.0.6 Materials availability

Not applicable

### 4.0.7 Code availability

Not applicable

### 4.0.8 Author contribution

SP and TA designed experiments. SP and DCF conducted experiments. SP analysed data and wrote the manuscript. SP, DCF and TA edited and proofread the manuscript.

## References

[1] Tang, P., Croxen, M.A., Hasan, M.R., Hsiao, W.W., Hoang, L.M.: Infection control in the new age of genomic epidemiology. American Journal of Infection Control 45(2), 170–179 (2017)

[2] Ten health issues WHO will tackle this year — who.int. https://www.who.int/news-room/spotlight/ten-threats-to-global-health-in-2019. x[Accessed 05-10-2024]

[3] Zhang, T., Zhang, X.-X., Ye, L.: Plasmid metagenome reveals high levels of antibiotic resistance genes and mobile genetic elements in activated sludge. PloS one 6(10), 26041 (2011)

[4] Murray, C.J., Ikuta, K.S., Sharara, F., Swetschinski, L., Aguilar, G.R., Gray, A., Han, C., Bisignano, C., Rao, P., Wool, E., et al.: Global burden of bacterial antimicrobial resistance in 2019: a systematic analysis. The lancet 399(10325), 629–655 (2022)

[5] Organization, W.H., et al.: Monitoring and evaluation of the global action plan on antimicrobial resistance: framework and recommended indicators (2019)

[6] Pillay, S., Calderón-Franco, D., Urhan, A., Abeel, T.: Metagenomic-based surveillance systems for antibiotic resistance in non-clinical settings. Frontiers in Microbiology 13, 1066995 (2022)

[7] Eyre, D.W.: Infection prevention and control insights from a decade of pathogen whole-genome sequencing. Journal of Hospital Infection 122, 180–186 (2022)

[8] Gerace, E., Mancuso, G., Midiri, A., Poidomani, S., Zummo, S., Biondo, C.: Recent advances in the use of molecular methods for the diagnosis of bacterial infections. Pathogens 11(6), 663 (2022)

[9] Bloemen, B., Gand, M., Vanneste, K., Marchal, K., Roosens, N.H., De Keersmaecker, S.C.: Development of a portable on-site applicable metagenomic data generation workflow for enhanced pathogen and antimicrobial resistance surveillance. Scientific Reports 13(1), 19656 (2023)

[10] Lee, M.H.-Y., Xu, G., Cheng, F., Khalid, A.F.: Testing surge capacity—a canadian covid-19 experience, ontario’s surge capacity for the first wave. Health Policy 125(10), 1291–1296 (2021)

[11] Loyola-Cruz, M.Á., Gonzalez-Avila, L.U., Martínez-Trejo, A., Saldaña-Padilla, A., Hernández-Cortez, C., Bello-López, J.M., Castro-Escarpulli, G.: Eskape and beyond: the burden of coinfections in the covid-19 pandemic. Pathogens 12(5), 743 (2023)

[12] Ramos-Mandujano, G., Grünberg, R., Zhang, Y., Bi, C., Guzmán-Vega, F.J., Shuaib, M., Gorchakov, R.V., Xu, J., Tehseen, M., Takahashi, M., et al.: An open-source, automated, and cost-effective platform for covid-19 diagnosis and rapid portable genomic surveillance using nanopore sequencing. Scientific Reports 13(1), 20349 (2023)

[13] Squitieri, L., Chung, K.C.: Surviving the covid-19 pandemic: surge capacity planning for nonemergent surgery. Plastic and Reconstructive Surgery 146(2), 437–446 (2020)

[14] Yingtaweesittikul, H., Ko, K., Abdul Rahman, N., Tan, S.Y.L., Nagarajan, N., Suphavilai, C.: Calmbelt: rapid sars-cov-2 genome characterization for outbreak tracking. Frontiers in medicine 8, 790662 (2021)

[15] Urhan, A., Abeel, T.: Emergence of novel sars-cov-2 variants in the netherlands. Scientific reports 11(1), 6625 (2021)

[16] Vigil, K., D’Souza, N., Bazner, J., Cedraz, F.M.-A., Fisch, S., Rose, J.B., Aw, T.G.: Long-term monitoring of sars-cov-2 variants in wastewater using a coordinated workflow of droplet digital pcr and nanopore sequencing. Water Research 254, 121338 (2024)

[17] Buenestado-Serrano, S., Herranz, M., Otero-Sobrino, Á., Molero-Salinas, A., Rodríguez-Grande, C., Sanz-Pérez, A., Galván, M.J.D., Catalán, P., Alonso, R., Muñoz, P., et al.: Accelerating sars-cov-2 genomic surveillance in a routine clinical setting with nanopore sequencing. International Journal of Medical Microbiology 314, 151599 (2024)

[18] Martin, C., Stebbins, B., Ajmani, A., Comendul, A., Hamner, S., Hasan, N.A., Colwell, R., Ford, T.: Nanopore-based metagenomics analysis reveals prevalence of mobile antibiotic and heavy metal resistome in wastewater. Ecotoxicology, 1–14 (2021)

[19] Serpa, P.H., Deng, X., Abdelghany, M., Crawford, E., Malcolm, K., Caldera, S., Fung, M., McGeever, A., Kalantar, K.L., Lyden, A., et al.: Metagenomic prediction of antimicrobial resistance in critically ill patients with lower respiratory tract infections. Genome medicine 14(1), 74 (2022)

[20] Freeman, C.N., Herman, E.K., Abi Younes, J., Ramsay, D.E., Erikson, N., Stothard, P., Links, M.G., Otto, S.J., Waldner, C.: Evaluating the potential of third generation metagenomic sequencing for the detection of brd pathogens and genetic determinants of antimicrobial resistance in chronically ill feedlot cattle. BMC veterinary research 18(1), 211 (2022)

[21] Białasek, M., Miłobędzka, A.: Revealing antimicrobial resistance in stormwater with minion. Chemosphere 258, 127392 (2020)

[22] Lee, A.W.-T., Ng, I.C.-F., Wong, E.Y.-K., Wong, I.T.-F., Sze, R.P.-P., Chan, K.-Y., So, T.-Y., Zhang, Z., Fung, S.K.-Y., Wong, S.C.-Y., et al.: Comprehensive identification of pathogenic microbes and antimicrobial resistance genes in food products using nanopore sequencing-based metagenomics. Food Microbiology 121, 104493 (2024)

[23] Bioworks, B.: Bento Lab: Experience Genetics Anywhere. 27 July 2018 (2018)

[24] Hirabayashi, A., Yanagisawa, H., Takahashi, H., Yahara, K., Boeing, P., Wolfenden, B., Nov, V., Lorn, V., Veng, M., Ann, V., et al.: On-site genomic epidemiological analysis of antimicrobial-resistant bacteria in cambodia with portable laboratory equipment. Frontiers in Microbiology 12, 675463 (2021)

[25] Marin, C., Marco-Jiménez, F., Martínez-Priego, L., De Marco-Romero, G., Soriano-Chirona, V., Lorenzo-Rebenaque, L., D’Auria, G.: Rapid oxford nanopore technologies minion sequencing workflow for campylobacter jejuni identification in broilers on site—a proof-of-concept study. Animals 12(16), 2065 (2022)

[26] Alcántara, R., Peñaranda, K., Mendoza-Rojas, G., Nakamoto, J.A., Martins-Luna, J., Valle-Mendoza, J., Adaui, V., Milón, P.: Unlocking sars-cov-2 detection in low-and middle-income countries. Cell reports methods 1(7) (2021)

[27] Ring, N., Low, A.S., Wee, B., Paterson, G.K., Nuttall, T., Gally, D., Mellanby, R., Fitzgerald, J.R.: Rapid metagenomic sequencing for diagnosis and antimicrobial sensitivity prediction of canine bacterial infections. Microbial Genomics 9(7), 001066 (2023)

[28] De Coster, W., D’hert, S., Schultz, D.T., Cruts, M., Van Broeckhoven, C.: Nanopack: visualizing and processing long-read sequencing data. Bioinformatics 34(15), 2666–2669 (2018)

[29] Portik, D.M., Brown, C.T., Pierce-Ward, N.T.: Evaluation of taxonomic classification and profiling methods for long-read shotgun metagenomic sequencing datasets. BMC bioinformatics 23(1), 541 (2022)

[30] Li, H.: Minimap2: pairwise alignment for nucleotide sequences. Bioinformatics 34(18), 3094–3100 (2018)

[31] Huson, D.H., Auch, A.F., Qi, J., Schuster, S.C.: Megan analysis of metagenomic data. Genome research 17(3), 377–386 (2007)

[32] Johnson, M., Zaretskaya, I., Raytselis, Y., Merezhuk, Y., McGinnis, S., Madden, T.L.: Ncbi blast: a better web interface. Nucleic acids research 36(suppl_2), 5–9 (2008)

[33] Alcock, B.P., Huynh, W., Chalil, R., Smith, K.W., Raphenya, A.R., Wlodarski, M.A., Edalatmand, A., Petkau, A., Syed, S.A., Tsang, K.K., et al.: Card 2023: expanded curation, support for machine learning, and resistome prediction at the comprehensive antibiotic resistance database. Nucleic acids research 51(D1), 690–699 (2023)

[34] Pellow, D., Mizrahi, I., Shamir, R.: Plasclass improves plasmid sequence classification. PLoS computational biology 16(4), 1007781 (2020)

[35] Gand, M., Bloemen, B., Vanneste, K., Roosens, N.H., De Keersmaecker, S.C.: Comparison of 6 dna extraction methods for isolation of high yield of high molecular weight dna suitable for shotgun metagenomics nanopore sequencing to detect bacteria. BMC genomics 24(1), 438 (2023)

[36] Bag, S., Saha, B., Mehta, O., Anbumani, D., Kumar, N., Dayal, M., Pant, A., Kumar, P., Saxena, S., Allin, K.H., et al.: An improved method for high quality metagenomics dna extraction from human and environmental samples. Scientific reports 6(1), 26775 (2016)

[37] Shamim, K., Sharma, J., Dubey, S.K.: Rapid and efficient method to extract metagenomic dna from estuarine sediments. 3 Biotech 7, 1–8 (2017)

[38] Lucena-Aguilar, G., Sánchez-López, A.M., Barberán-Aceituno, C., Carrillo-Avila, J.A., López-Guerrero, J.A., Aguilar-Quesada, R.: Dna source selection for down-stream applications based on dna quality indicators analysis. Biopreservation and biobanking 14(4), 264–270 (2016)

[39] Koetsier, G., Cantor, E.: A practical guide to analyzing nucleic acid concentration and purity with microvolume spectrophotometers. New England Biolabs Inc, 1–8 (2019)

[40] Technologies, O.N.: Contaminants. https://community.nanoporetech.com/contaminants Accessed 2024

[41] Technologies, O.N.: gDNA. https://community.nanoporetech.com/extraction_method_groups/optional-fragmentation-of-gdnas Accessed 2024

[42] Maguire, M., Kase, J.A., Brown, E.W., Allard, M.W., Musser, S.M., González-Escalona, N.: Metagenomic survey of agricultural water using long read sequencing: considerations for a successful analysis. Frontiers in Environmental Science 10, 830300 (2022)

[43] Technologies, O.N.: Flongle. https://nanoporetech.com/products/sequence/flongle Accessed 2024

[44] Berbers, B., Ceyssens, P.-J., Bogaerts, P., Vanneste, K., Roosens, N.H., Marchal, K., De Keersmaecker, S.C.: Development of an ngs-based workflow for improved monitoring of circulating plasmids in support of risk assessment of antimicrobial resistance gene dissemination. Antibiotics 9(8), 503 (2020)

[45] Gehrig, J.L., Portik, D.M., Driscoll, M.D., Jackson, E., Chakraborty, S., Gratalo, D., Ashby, M., Valladares, R.: Finding the right fit: evaluation of short-read and long-read sequencing approaches to maximize the utility of clinical microbiome data. Microbial Genomics 8(3), 000794 (2022)

[46] Li, S., Mann, D.A., Zhang, S., Qi, Y., Meinersmann, R.J., Deng, X.: Microbiome-informed food safety and quality: longitudinal consistency and cross-sectional distinctiveness of retail chicken breast microbiomes. Msystems 5(5), 10–1128 (2020)

[47] Sequino, G., Cobo-Diaz, J.F., Valentino, V., Tassou, C., Volpe, S., Torrieri, E., Nychas, G.-J., Ordóñez, A.Á., Ercolini, D., De Filippis, F.: Microbiome mapping in beef processing reveals safety-relevant variations in microbial diversity and genomic features. Food Research International, 114318 (2024)

[48] Spieck, E., Hartwig, C., McCormack, I., Maixner, F., Wagner, M., Lipski, A., Daims, H.: Selective enrichment and molecular characterization of a previously uncultured nitrospira-like bacterium from activated sludge. Environmental Microbiology 8(3), 405–415 (2006)

[49] Niestępski, S., Harnisz, M., Ciesielski, S., Korzeniewska, E., Osińska, A.: Environmental fate of bacteroidetes, with particular emphasis on bacteroides fragilis group bacteria and their specific antibiotic resistance genes, in activated sludge wastewater treatment plants. Journal of hazardous materials 394, 122544 (2020)

[50] Kowal, P., Ciesielski, S., Godzieba, M., Fitobór, K., Gajewska, M., Kołecka, K.: Assessment of diversity and composition of bacterial community in sludge treatment reed bed systems. Science of The Total Environment 756, 144060 (2021)

[51] Wrenn, D.C., Drown, D.M.: Nanopore adaptive sampling enriches for antimicrobial resistance genes in microbial communities. GigaByte 2023 (2023)

[52] Venkateswaran, P., Vasudevan, S., David, H., Shaktivel, A., Shanmugam, K., Neelakantan, P., Solomon, A.P.: Revisiting eskape pathogens: Virulence, resistance, and combating strategies focusing on quorum sensing. Frontiers in cellular and infection microbiology 13, 1159798 (2023)

[53] Kelly, J.B., Nolan, A.C., Zeden, M.S.: How can we escape the eskapes: Antimicrobial resistance mechanisms and what lies ahead? PLoS pathogens 20(6), 1012270 (2024)

[54] Savin, M., Bierbaum, G., Hammerl, J.A., Heinemann, C., Parcina, M., Sib, E., Voigt, A., Kreyenschmidt, J.: Eskape bacteria and extended-spectrum-β-lactamase-producing escherichia coli isolated from wastewater and process water from german poultry slaughterhouses. Applied and Environmental Microbiology 86(8), 02748–19 (2020)

[55] Russo, T.P., Minichino, A., Gargiulo, A., Varriale, L., Borrelli, L., Pace, A., Santaniello, A., Pompameo, M., Fioretti, A., Dipineto, L.: Prevalence and phenotypic antimicrobial resistance among eskape bacteria and enterobacterales strains in wild birds. Antibiotics 11(12), 1825 (2022)

[56] Wang, L., Tang, J.-W., Li, F., Usman, M., Wu, C.-Y., Liu, Q.-H., Kang, H.-Q., Liu, W., Gu, B.: Identification of bacterial pathogens at genus and species levels through combination of raman spectrometry and deep-learning algorithms. Microbiology spectrum 10(6), 02580–22 (2022)

[57] Dijk, L.R., Walker, B.J., Straub, T.J., Worby, C.J., Grote, A., Schreiber IV, H.L., Anyansi, C., Pickering, A.J., Hultgren, S.J., Manson, A.L., et al.: Strainge:b a toolkit to track and characterize low-abundance strains in complex microbial communities. Genome biology 23(1), 74 (2022)

[58] Heir, E., Moen, B., Åsli, A.W., Sunde, M., Langsrud, S.: Antibiotic resistance and phylogeny of pseudomonas spp. isolated over three decades from chicken meat in the norwegian food chain. Microorganisms 9(2), 207 (2021)

[59] Wang, W., Peng, Z., Baloch, Z., Hu, Y., Xu, J., Zhang, W., Fanning, S., Li, F.: Genomic characterization of an extensively-drug resistance salmonella enterica serotype indiana strain harboring blandm-1 gene isolated from a chicken carcass in china. Microbiological research 204, 48–54 (2017)

[60] Sun, Y., Hu, X., Guo, D., Shi, C., Zhang, C., Peng, X., Yang, H., Xia, X.: Disinfectant resistance profiles and biofilm formation capacity of escherichia coli isolated from retail chicken. Microbial Drug Resistance 25(5), 703–711 (2019)

[61] Vieira, T.R., Oliveira, E.F.C., Cibulski, S.P., Silva, N.M., Borba, M.R., Oliveira, C.J., Cardoso, M.: Comparative resistome, mobilome, and microbial composition of retail chicken originated from conventional, organic, and antibiotic-free production systems. Poultry Science 102(11), 103002 (2023)

[62] Filipic, B., Novovic, K., Studholme, D.J., Malesevic, M., Mirkovic, N., Kojic, M., Jovcic, B.: Shotgun metagenomics reveals differences in antibiotic resistance genes among bacterial communities in western balkans glacial lakes sediments. Journal of water and health 18(3), 383–397 (2020)

[63] Anes, J., McCusker, M.P., Fanning, S., Martins, M.: The ins and outs of rnd efflux pumps in escherichia coli. Frontiers in microbiology 6, 587 (2015)

[64] Chauviat, A., Meyer, T., Favre-Bonté, S.: Versatility of stenotrophomonas maltophilia: Ecological roles of rnd efflux pumps. Heliyon 9(4) (2023)

[65] Ni, R.T., Onishi, M., Mizusawa, M., Kitagawa, R., Kishino, T., Matsubara, F., Tsuchiya, T., Kuroda, T., Ogawa, W.: The role of rnd-type efflux pumps in multidrug-resistant mutants of klebsiella pneumoniae. Scientific reports 10(1), 10876 (2020)

[66] Cooper, A.L., Low, A., Wong, A., Tamber, S., Blais, B.W., Carrillo, C.D.: Modeling the limits of detection for antimicrobial resistance genes in agri-food samples: a comparative analysis of bioinformatics tools. BMC microbiology 24(1), 31 (2024)

[67] Petersen, L.M., Martin, I.W., Moschetti, W.E., Kershaw, C.M., Tsongalis, G.J.: Third-generation sequencing in the clinical laboratory: exploring the advantages and challenges of nanopore sequencing. Journal of clinical microbiology 58(1), 10–1128 (2019)

[68] Smalla, K., Jechalke, S., Top, E.M.: Plasmid detection, characterization, and ecology. Plasmids: biology and impact in biotechnology and discovery, 445–458 (2015)

